# Evaluation of recombination rate of microhaplotypes based on the 1000 Genomes Project family data

**DOI:** 10.1101/2023.12.11.571057

**Authors:** Yifan Wei, Xi Li, Tiantian Shan, Haoyu Wang, Xuan Dai, Qiang Zhu, Yufang Wang, Ji Zhang

## Abstract

Microhaplotype, a kind of multi-allelic genetic marker that contains two or more SNPs in the range of hundreds of base pairs, has emerged as an important genetic marker in forensic genetics. However, the recombination within the microhaplotype, which has been neglected in most studies, leads to changes in its alleles during inheritance, producing alleles that are not identical to parental alleles and affecting the likelihood ratio calculation of the MH in forensic DNA analysis involving kinship. We constructed a highly dense and extensive MH set in the human genome based on the expanded 1000 Genomes Project (GRCh38). Using the pedigree data contained in 18 populations and a total of 1204 meioses in the GRCh38 dataset, we scanned and confirmed the recombination events that occurred within the above MH loci, to clarify the relationship between recombination and MH polymorphisms. We investigated the average incidence of MH recombinant alleles and the average recombination rate of genome-wide MHs in the hope of providing a reference for the correction of the likelihood ratio in the MH system of future kinship analysis.

## 1. Introduction

Microhaplotype (MH), a kind of multi-allelic genetic marker that contains two or more SNPs in the range of hundreds of base pairs [1], has received considerable critical attention from forensic geneticists. Since its widespread distribution in the whole genome [2], high polymorphism, and specific distribution in the population [3] to a certain extent, microhaplotype can be well applied to individual identification [4–6] and ancestry inference [7–9]. For kinship analysis, whereas, microhaplotypes should be applied with caution, as mutations or recombinations between SNPs within the MH may result in offspring alleles that are not identical to the parental alleles and thus affect the analysis of kinship.

The 1000 Genomes Project has performed deep sequencing of family samples to directly estimate *de novo* germline base mutation [10], about 10^-8^. For the mutation rate of MH, since SNPs are components of MH, their average mutation rate is thought to be low and negligible. However, there is still a lack of proper assessment of the recombination issue between SNPs within microhaplotypes. To avoid the effects of recombination, researchers selected MHs located in regions with an average recombination rate of less than 1% per megabase or not in recombination hotspots [11]. On the premise of this average recombination rate, some scholars [12] believe that the probability of recombination within a few hundred bases of MH is about 10^−6^ or even lower, and the impact on the relationship analysis can be ignored. Nevertheless, the recombination rate within MHs estimated in this way remains to be verified because the calculation of 1 cM/Mb is not entirely based on pedigree data and observation of recombination events in meiosis. It is imperative to study its recombination rate, thereby further refining the screening criteria.

There have been a variety of methods to be used for the study of human genetic recombination so far. On the basis of coalescent theory and population genetics, the HapMap project made use of composite likelihood to develop the most thorough genetic map [13], but differential information on recombination rate to individuals or gender was not available, and the computation cost considerably. The recombination could be identified by family samples. This method phased the genetic markers by linkage analysis and directly observed recombination in the gametes, yet the resolution of the genetic map was limited by relatively few meioses and low-density markers. The resolution reached 10–70 kb merely though high-density SNPs were used later [14–16]. In recent years, some researchers have also used the method of single sperm sequencing to observe the occurrence of recombination in meiosis with a resolution of up to 250 bp [17].

The expanded 1000 Genomes Project (e1kGP) was published in 2022 [18]. In contrast to 1000 Genomes Project (1kGP) phase 3, e1kGP has significantly improved sensitivity and specificity for calling SNVs with a lower false discovery rate (FDR) and higher quality of haplotype phase. Hence, we constructed an MH set based on e1kGP sequencing data and analyzed all 602 trios to discern recombination events in the range of 350 bp (i.e., within MHs) by observing the 1204 meioses. We investigated the correlation between MH polymorphism and recombination, as well as the recombination rate within MHs.

## 2. Materials and Methods

### 2.1. Sample preparation

Phased whole genome sequencing (WGS) data of 2504 unrelated individuals who were the same individuals with 1kGP phase3 from 26 populations were downloaded from the expanded 1000 Genomes Project (e1kGP) (http://ftp.1000genomes.ebi.ac.uk/vol1/ftp/data_collections/1000G_2504_high_coverage/working/20201028_3202_phased/). These unrelated individuals from 26 populations were used to screen SNPs and construct MHs, thereby calculating forensic parameters. 602 trios from 18 populations were used to observe recombination (Supplemental Table S1) [18].

### 2.2. Preliminary screening of SNPs

Preliminary screening of SNPs was performed by the shell script based on UNIX and VCFtools according to two criteria below: 1) All SNPs were located in 1–22 autosomes, and 2) conformed to Hardy-Weinberg equilibrium (p > 0.05). Note that we did not limit the minor allelic frequency (MAF) of SNP to ensure better compatibility between different populations for the constitution of MHs.

### 2.3. Construction of microhaplotypes

Using the screened SNPs as input, we developed a Python script that could exhaust all possibilities of SNP combination and construct an MH set on the basis of the segment length of MH specified, that is, 350 bp in this study. The specific implementation process was as follows (Fig. 1): First, the SNP at the 5’ end (the numerically lowest physical position) of each chromosome was used as the start, and the adjacent SNPs were added one by one downstream to construct the MHs in the range of 350 bp; Once the length exceeded 350 bp, move the coordinate of the “start” SNP downstream one position (i.e., one SNP) and repeat the steps above. The process did not terminate until the 3’ end of the chromosome. Second, we dropped the subset-MHs, whose starting points and terminal points were both included in the longest MHs. In this step, we still kept the intersection-MHs, one end of which was inside the longest MH and one end outside the longest MH. Third, we discarded the intersection-MHs and only retained the nonoverlapping MHs, whose starting points were after the terminal points of the previous MHs. After searching across all 22 autosomes, we obtained the MH set. We anchored SNP physical positions to guarantee that different populations had the same MH loci, meaning the same physical lengths and positions of MHs in each population, in this case, some SNPs in the MHs of the populations might not have polymorphism. The fundamental information of all MHs among 18 populations was counted and calculated by our in-house Python scripts, including the segment length (bp), the number of SNPs constituting each MH, observed heterozygosity (Ho), discrimination power (DP), and the effective number of alleles (Ae) according to unrelated individuals from these populations.

**Figure 1.**
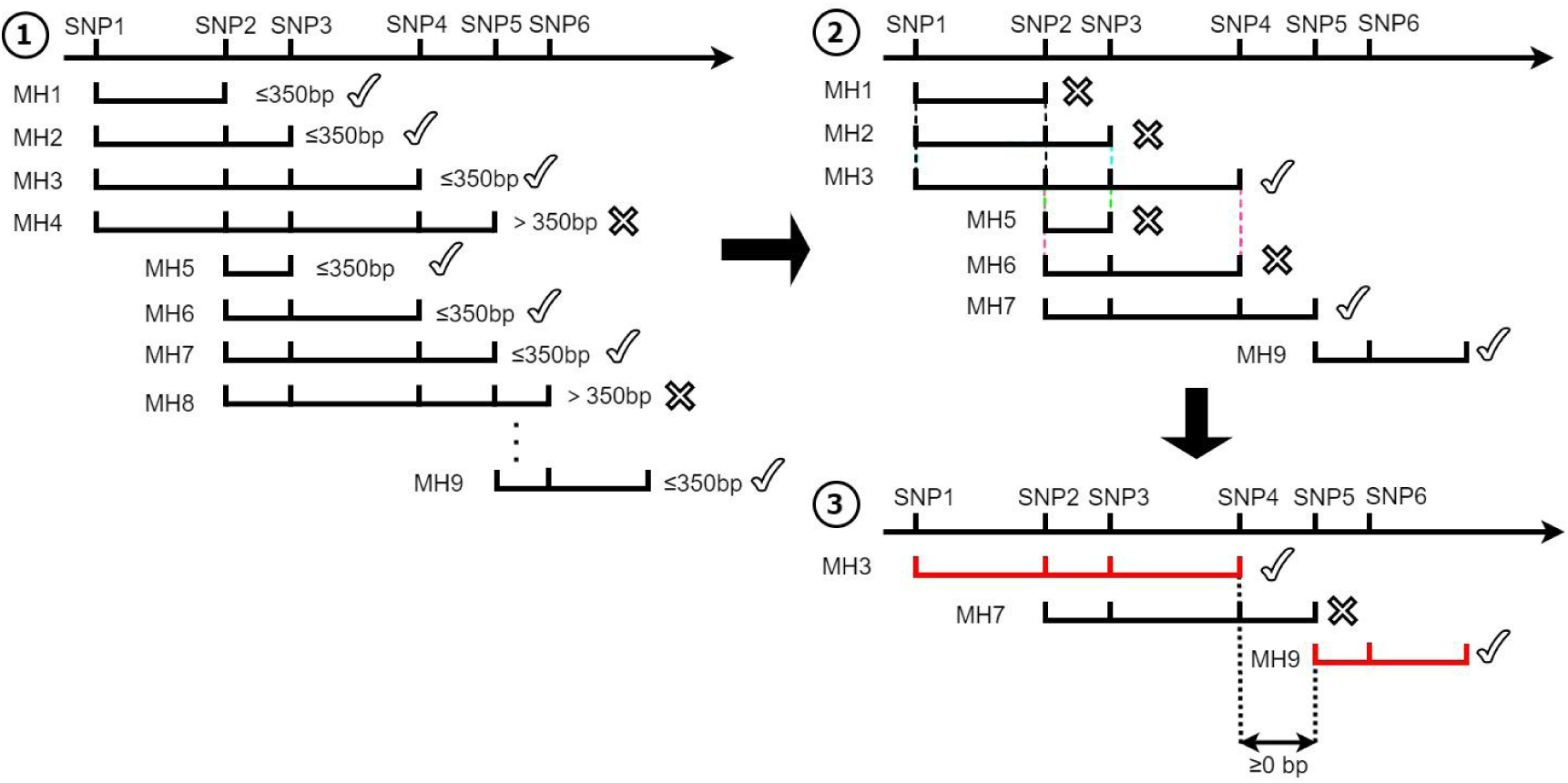
Assembly process of MHs. Step 1: SNP1 was used as the start, and adjacent SNP was added one by one backward to construct the MHs until the length exceeded 350 bp. Move the coordinate of the “start” SNP to the second SNP (SNP2) and repeat the steps above. The process did not terminate until the 3’ end of the chromosome. Step 2: the subset-MHs (MH1, MH2, MH3, MH5, MH6), whose starting points and terminal points were both included in the longest MHs, were dropped (marked as crosses). In this step, we still kept the intersection-MHs (MH7 and MH3 had an intersection), one end of which was inside the longest MH and one end was outside the longest MH. Step 3: we discarded the intersection-MHs (MH7 in this example) and only retained the nonoverlapping MHs (marked as ticks and red solid lines, i.e., MH3 and MH9 in the figure), whose starting points were after the terminal points of the previous MHs.

### 2.4. Identification of variant alleles

Using WGS data of 602 trios from 18 populations, we developed Python scripts identifying MH variant alleles. First, all MH alleles of trios were extracted. Then, MH alleles were compared between offspring and parents. If any of offspring alleles did not match parental alleles, the genotypes of the trio were output to discriminate the cause of new alleles subsequently.

### 2.5. Determining the position and type of the variation

For convenience, four alleles of a father and a mother on a certain MH were referred to as F1, F2, and M1, M2, respectively, from which two alleles of an offspring were referred to as C1, and C2 through meiosis. The positions and type of the variation could be identified by comparing the alleles of offspring and parents on the condition that MH alleles were phased and parental origin was known. For example, assuming that C1 was inherited from F1 (F1→C1) and C2 from M1 (M1→C2), C1, and C2 should be the same with F1, M1, respectively, on the basis of Mendel’s laws of inheritance. However, F2 would be considered additionally to determine the position and the cause of allele variation when C1 was not the same as F1. When we had compared every single SNP between F1 and C1 following a 5’ to 3’ direction, with the first inconsistent base identified and the same positional base in F2 being consistent with C1 in the meantime, we regarded this situation as the recombination and skipped from the base of F1 to F2 to search backward continually. Otherwise, the base was marked as a mutation or a sequencing error (Fig. 2A). Meanwhile, only heterozygous SNPs within the MH locus could reveal whether the alleles are recombinant, indicating that the position of recombination could be located in the region spanned by the two closest heterozygous SNPs (also referred to as SNP interval). There were three kinds of situations (Fig. 2).

**Figure 2.**
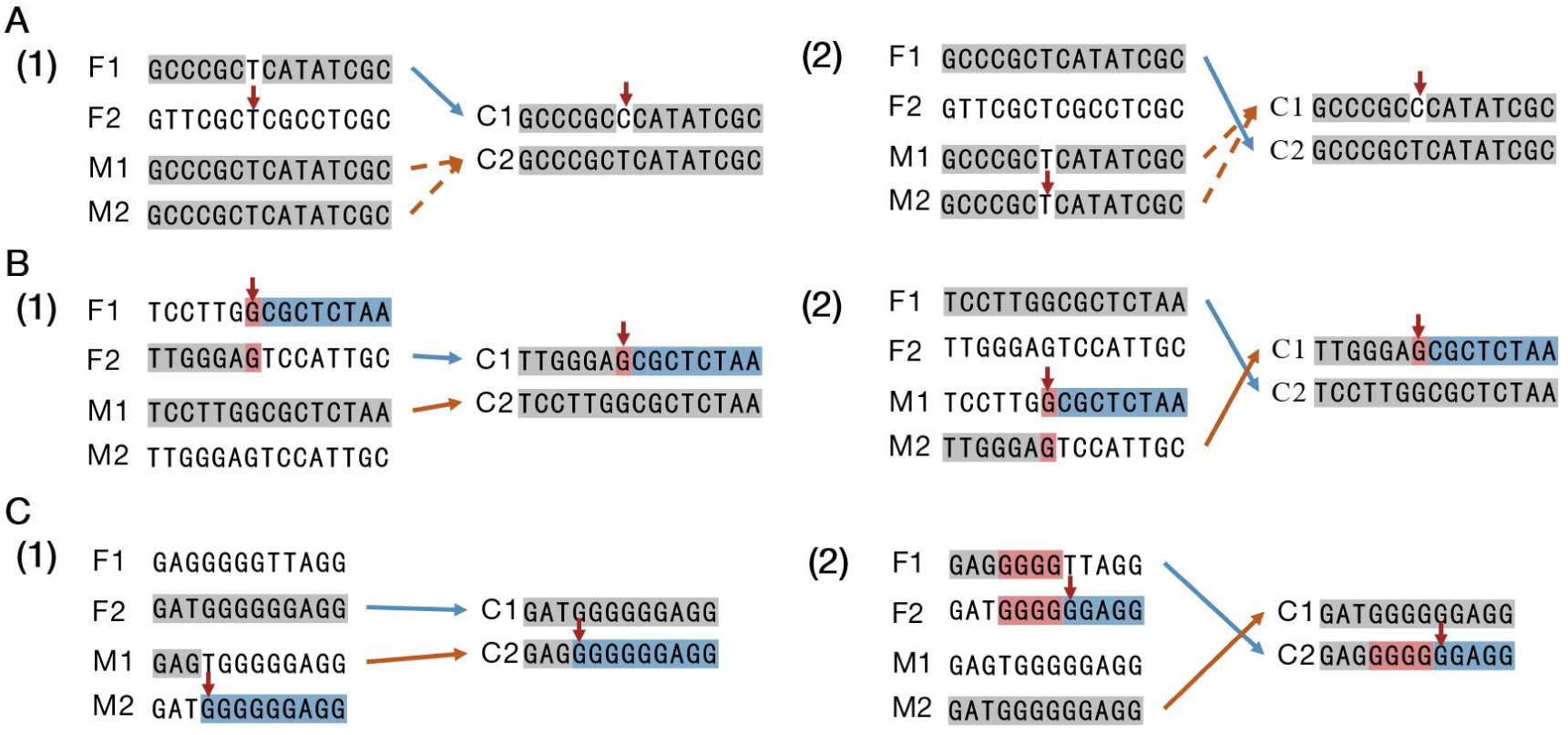
Determining the position and type of variation of MH alleles. In the trios, the father’s alleles are F1 and F2, and mother’s alleles are M1 and M2, and the offspring’s alleles are C1 and C2. Assume the parental origin is F1 → C1 and M1 → C2. A: (1) Allele C2 is from M1 or M2, and C1 originates from sequencing error or mutation. (2) Allele C2 is from F1 or F2, and C1 originates from sequencing error or mutation. B: F1 and F2 (1), or M1 and M2 (2) undergo recombination to produce C1, and the recombination position is the 7th SNP (highlighted in red). C: The recombination intervals are not identical according to the different parental origins, and these kinds of alleles are classified as variant alleles with undetermined recombination locations.

Every possible parental origin, however, needed to be considered in that whether each of the offspring alleles was paternal or maternal was unknown. Taking into account every possibility, if variant alleles could only be explained by mutation or sequencing error, they were regarded as “non-recombinant alleles”. Otherwise, as few recombination events as possible were retained, since the segment length of all MH loci was less than 350 bp and the possibility of multiple recombination events was extremely low at such a fine resolution. Besides, among such MH alleles for which the cause of variation could be identified as SNP recombination, there was still a fraction for which the specific parental origin could not be clarified, because the combination of gametes in which the minimum number of recombination occurred was not unique. For instance, when parents had identical genotypes, offspring alleles failed to be phased but were still output as “recombinant alleles” on account of the inference of a clear recombination interval. If recombination events inferred from various parental origins occurred within different SNP intervals, these alleles were classified as “variant alleles with undetermined recombination location”. On the basis of the analysis principles above (Fig. 3), we developed a set of Python scripts and identified and analyzed MH variant alleles acquired by the previous section automatically to deduce the source, position, and three types of variations. Output was sampled randomly to be validated manually to guarantee the accuracy of scripts.

**Figure 3.**
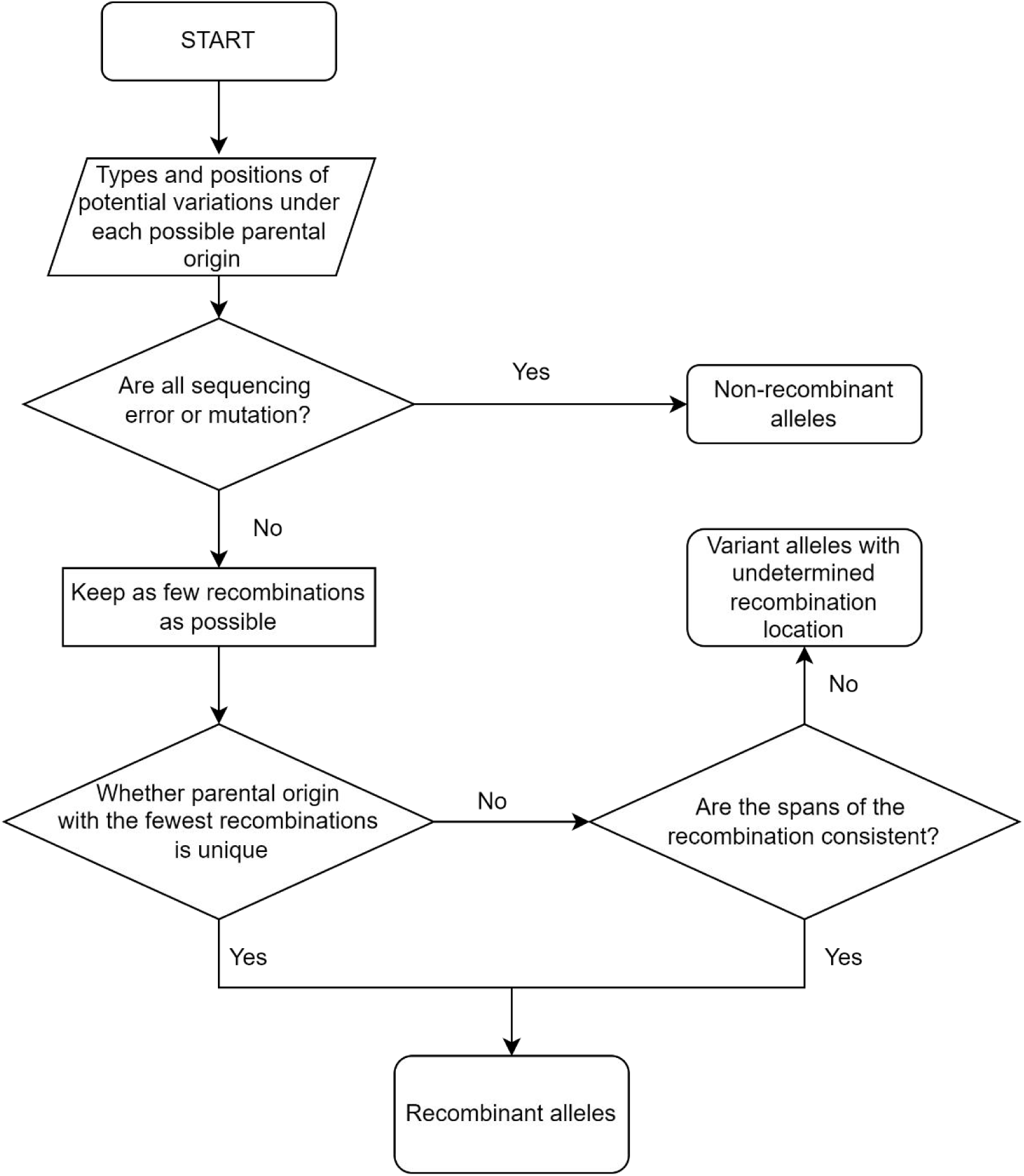
The flow chart of inferring the position and type of variation of MH alleles.

### 2.6. Data analysis

As mentioned above, our study focused on recombination within MH and estimated the effect of recombination on genetic variations conservatively. Hence, among the three situations, recombinant alleles, whose recombinant intervals were explicit, were retained to be analyzed and calculated while the other two kinds of alleles were discarded and not analyzed further.

For the sake of understanding the relation intuitively between recombination within MH and MH polymorphism, a series of parameters were introduced and calculated. The proportion of recombinant MH loci (Pro_r) was the ratio of the number of recombinant MH loci and the total number of MH loci. Recombinant MH loci were the MH loci where “new” and “recombinant alleles” were generated.

To measure the likelihood of recombinant alleles arising during the transmission of MH from parents to offspring, we calculated the incidence of recombinant alleles, 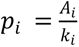(*i* = *1*,*2*,…,*n*), where *k_i_* was the total number of gamete alleles and *A_i_* was the number of recombinant alleles. With *n* MH loci tested, the average incidence of recombinant alleles of these MHs (AvgP) was calculated as below:

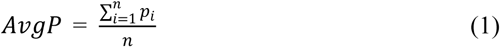

For autosomal loci, the value of *k_i_* was double the number of trios observed and equaled to *m*, the number of meiosis observed. Thus, we obtained:

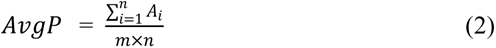

The recombination rate r within each MH was defined as 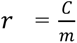, where *C* was the number of recombination observed between SNPs constructing one MH, and *m* was the number of meiosis observed. Note that the recombination rate in this research refers to the probability of occurring recombination between SNPs within MHs. Assuming that there were *n* MH loci, and *r_i_* (i=1, 2, …, n) was the recombination rate of each MH locus, the average recombination rate (AvgRR) of all loci was calculated as below:

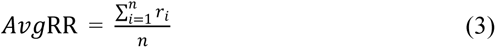

Thus, we could get:

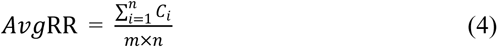

*C_i_* was the number of recombinations observed within MH locus i (i=1, 2, …, n). The average recombination rate (AvgRR) of MHs was similar to the average mutation rate of STRs.

In addition, with the human genome recombination hotspots and the genetic map published by International HapMap Project as a reference, the referenced recombination rate of MHs (ld_based_rate) based upon HapMap was calculated and compared with the average recombination rate of MHs in our study. Considering that the MH dataset in this study was based on the human reference genome GRCh38, UCSC liftOver (https://genome.ucsc.edu/cgi-bin/hgLiftOver) was employed to convert physical coordinates from HapMap to GRCh38 after which SNPs not consistent between them were removed. The referenced average rate of recombination was calculated below:

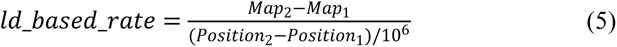

*Map*_1_ and *Map*_2_ were genetic distances of the SNPs with the closest distances upstream and downstream of the target site in the HapMap genetic map, respectively. Similarly, *Position*_1_ and *Position*_2_ were physical distances of the SNPs.

### 2.7. High-density distribution regions

According to the distribution of recombinant MHs in the genome, there were some visible high-density regions observed. To avoid the effects taken by these regions and access the recombination situation as conservatively as possible, we removed them and extended the deleted regions by 5 Mb each forward and backward. After that, we recalculated the three measures (Pro_r, AvgP, AvgRR).

## 3. Results

### 3.1. Constructing microhaplotypes

After extracting and filtering VCF files of 2504 unrelated individuals from e1KGP, we obtained 47,875,562 SNPs (Supplemental Table S2) to construct MHs while the average density was 16.65 SNPs/1 kb.

With the longest length of MH set as 350 bp, the MH dataset including 6,983,511 MH loci was obtained for the observation of meiosis. The number of MHs on each chromosome ranged from 90,231 to 619,027 (Supplemental Table S2). On average, there were 2,429 MHs per Mb (6,983,511/2,875 Mb) in the human reference genome. From the view of distribution density, 1,817 MHs per Mb were detected on chromosome 22, which was the sparsest, and 2,576 MHs per Mb were detected on chromosome 3, which was the densest. The physical positions of MHs in the human genome illustrated the overall uniform distribution, except for the low-density regions around the gaps of GRCh38 and some known highly variable regions such as MHC (Fig. 4A).

**Figure 4.**
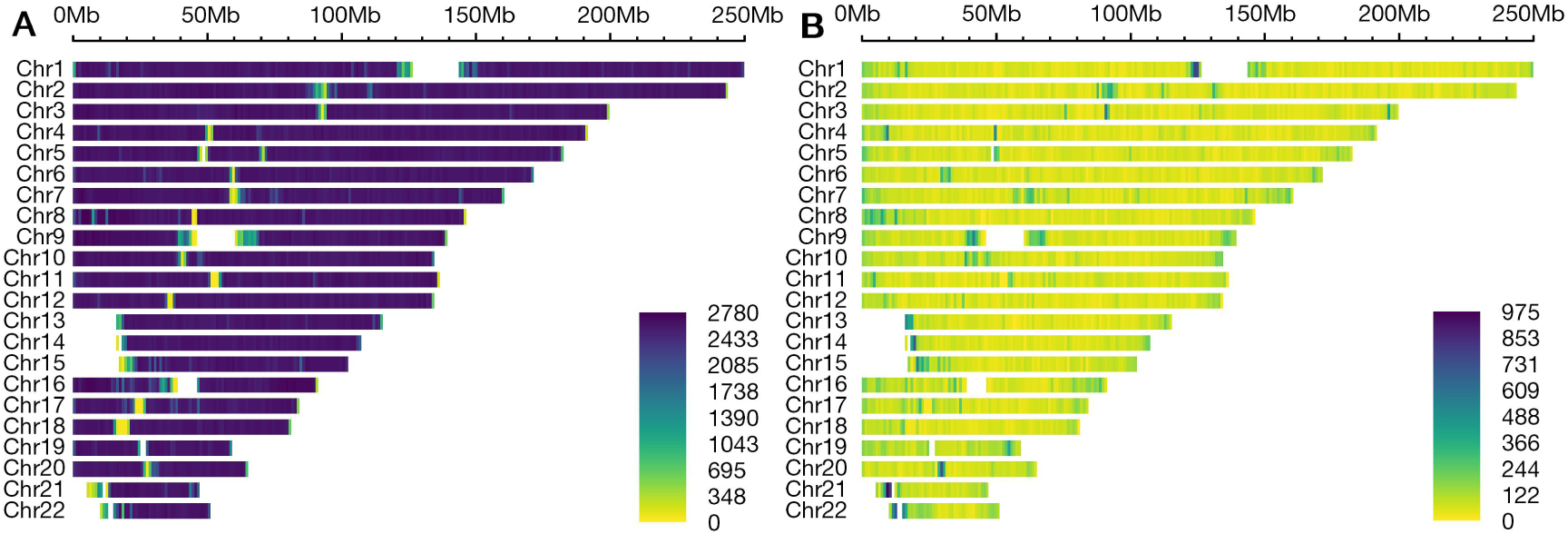
Distribution of microhaplotype loci and recombinant MHs in the human genome. Refer to GRCh38. Color scale shades indicate the number of MHs per million base pairs (Mb). A: All MHs in the genome. The figure was plotted using the MH dataset, totaling 6,983,511 MHs, excluding the sex chromosome dataset. B: All recombinant MHs of 18 populations. Note that each population has different recombinant MHs, and we plotted the figure using the union of 18 sets.

Of the all 2,875 Mb in 22 autosomes (http://hgdownload.soe.ucsc.edu/goldenPath/hg38/bigZips/hg38.chrom.sizes), 2,220 Mb were contained in the MH dataset of our MH library by which the overall coverage rate was 77.21% while the coverage rate of each chromosome ranged from 58.06% (Chr 22) to 81.76% (Chr 4). The number of SNPs of each MH locus was an average of 7.38 SNPs (range from 2 to 310 SNPs). Besides, the segment length of MHs ranged from 2 bp to 350 bp, with a mean of 318 bp.

The effective number of alleles (Ae) is calculated as 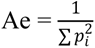, where *p_i_* is the frequency of allele *i* [19], which is used to measure the variation level of one MH within a specific population. Among 18 populations, the number of MHs decreased with the Ae value increasing (Supplemental Table S3). The top 3 populations with the highest Ae were IBS, STU, and ACB, the values of which were 27.72, 27.67, and 26.83, respectively.

### 3.2. Variant alleles induced by recombination

575,752 variant alleles were detected from the offspring’s alleles of the 18 populations. We reserved 451,442 recombinant alleles whose recombination positions could be confirmed explicitly, accounting for 78.41%, while the rest were not recombinant alleles or recombinant alleles whose recombinant positions could not be confirmed. GWD had the highest percentage of 16.01% (72,287), with ITU accounting for the lowest 0.35% (1,573) (Fig. 5). The top 3 populations with the highest number of recombinant alleles were GWD, YRI, and ESN, concentrating in the African continent.

**Figure 5.**
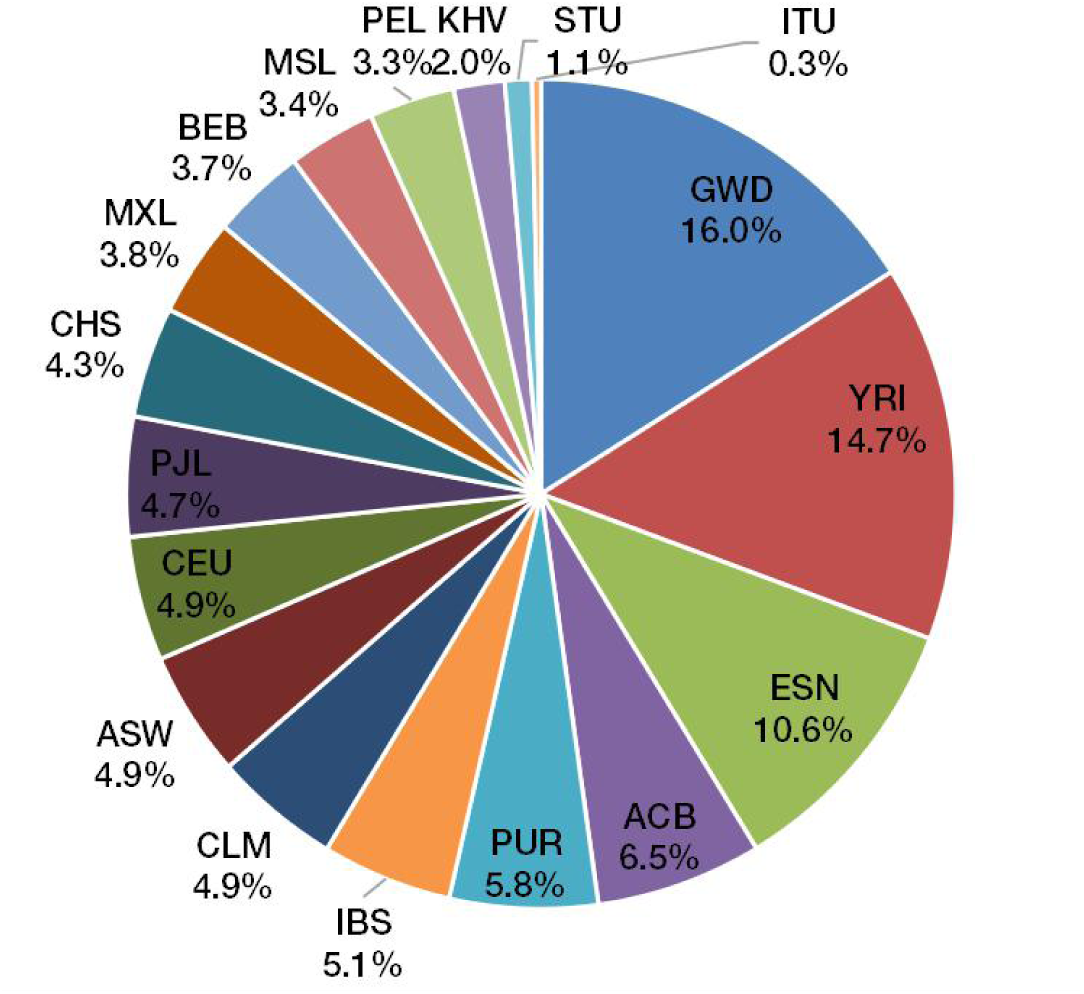
The percentage of recombinant alleles of each population.

### 3.3. Recombinant microhaplotypes

We accordingly counted the number of MH loci for which recombinant alleles were observed and presented their distribution in a total of 18 populations (hereafter referred to as 18-sum population) (Fig. 4B). Especially, GWD contained the most recombinant loci (48,457), accounting for 22.5% among all recombinant MHs. The number of recombinant MHs in YRI and ESN populations ranked after GWD. We counted the number of recombinant MHs shared between populations and unique to each population (Fig. 6), showing that all 18 populations shared only 4 loci (Fig. 6B). These four loci were located near the MHC of chromosome 6 and near the proximal telomeres of chromosomes 13 and 14 (Table 1). Considering that some populations possessed relatively few trios that probably masked the features of shared loci, we removed the populations with fewer than 30 trios and retained 12 populations to reassess the number of shared loci. The 12 populations shared 126 recombinant MHs, most of which were located near the MHC, the proximal telomeres, and centromeres (Fig. 6C, D). Among 18 populations, the number of unique loci of GWD was 24,618, which was the most (Fig. 6A). Further, we counted recombinant MHs with different numbers of recombinant alleles for the 18-sum population. Overall, each recombinant MH possessed between 1 and 144 recombinant alleles among 18 populations, with most loci (74.95%) having only one recombinant allele (Table 2). The MH locus CHR13_000009856, which recombined 223 times in 135 trios and generated up to 144 recombinant alleles (Table 2), was one of four loci shared by 18 populations.

**Figure 6.**
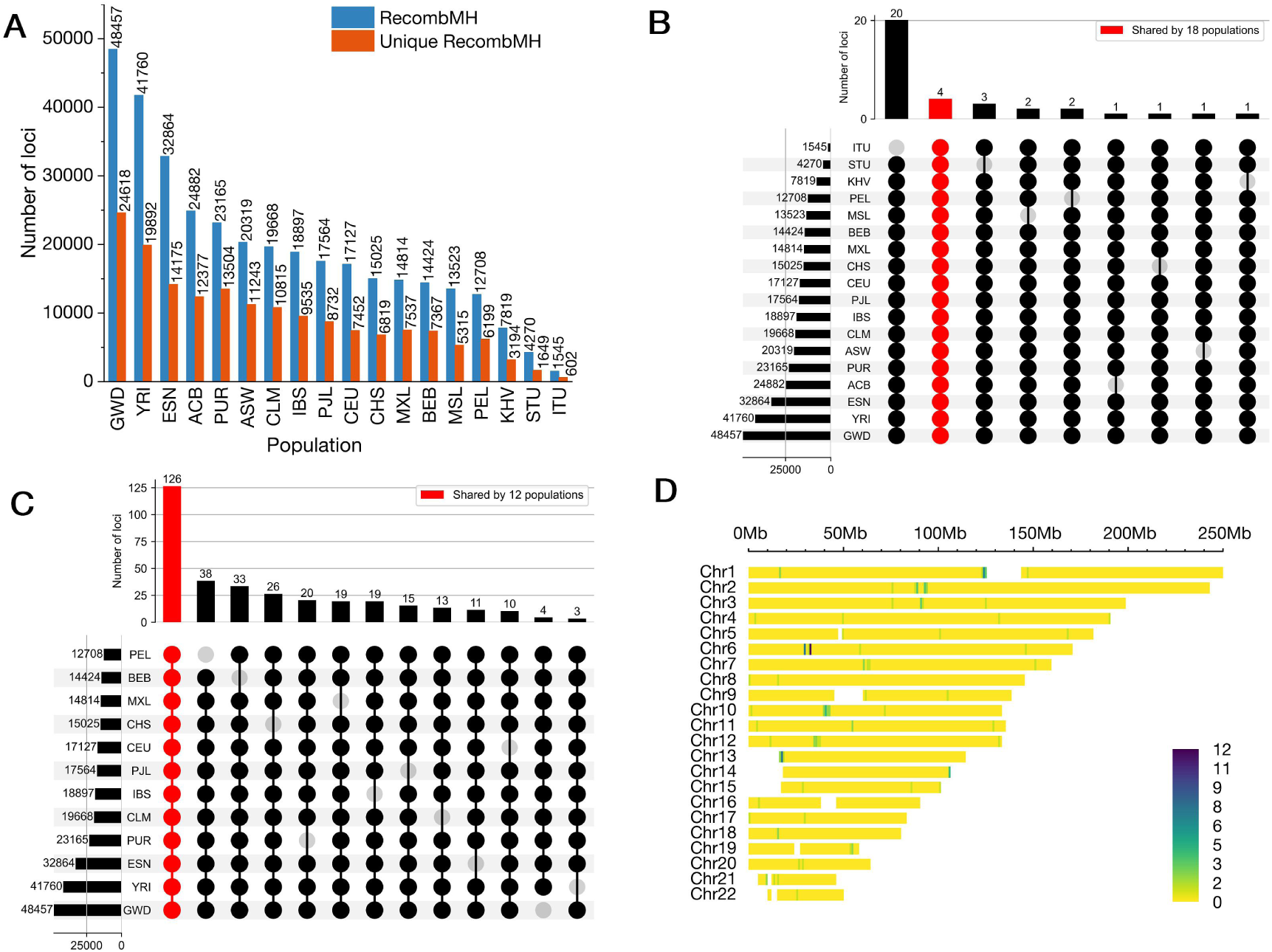
The number of recombinant MHs of 18 populations and the number of shared loci between them. B and C show the cardinality of different combinations. A: The number of unique recombinant MHs of each population. Blue bars indicate recombinant MHs, with orange bars indicating unique recombinant MHs in each population. B: All columns except for the second one count loci in each combination of exactly 17-population sets, with the second column (the red one) counting loci presenting in all 18 populations. C: 126 MHs are shared by 12 populations (red column), each of which has more than 30 trios. The rest columns show the number of loci shared by 11 populations. D: Distribution of 126 MHs shared by 12 populations on the genome. Darker color indicates a higher number of loci.

**Table 1.**
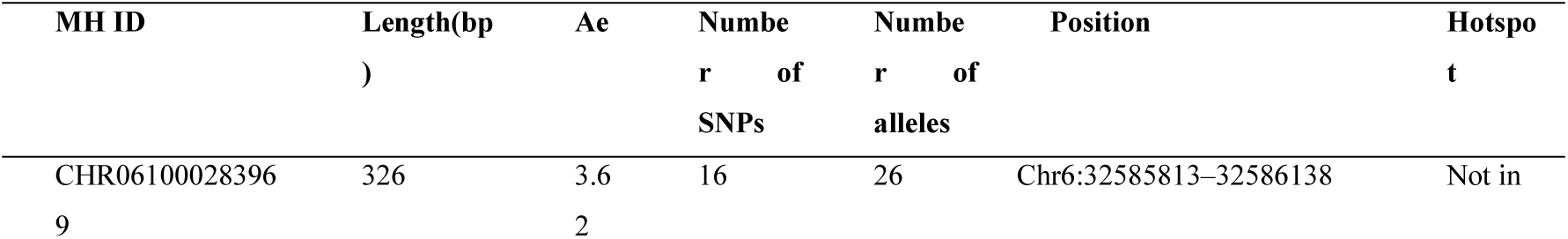
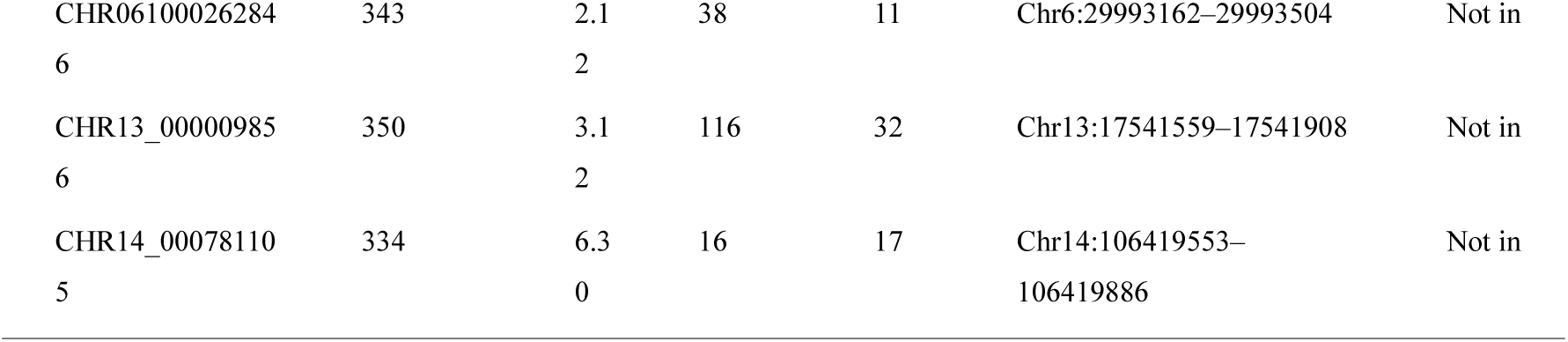
Information of 4 shared recombinant MHs by 18 populations.

**Table 2.**
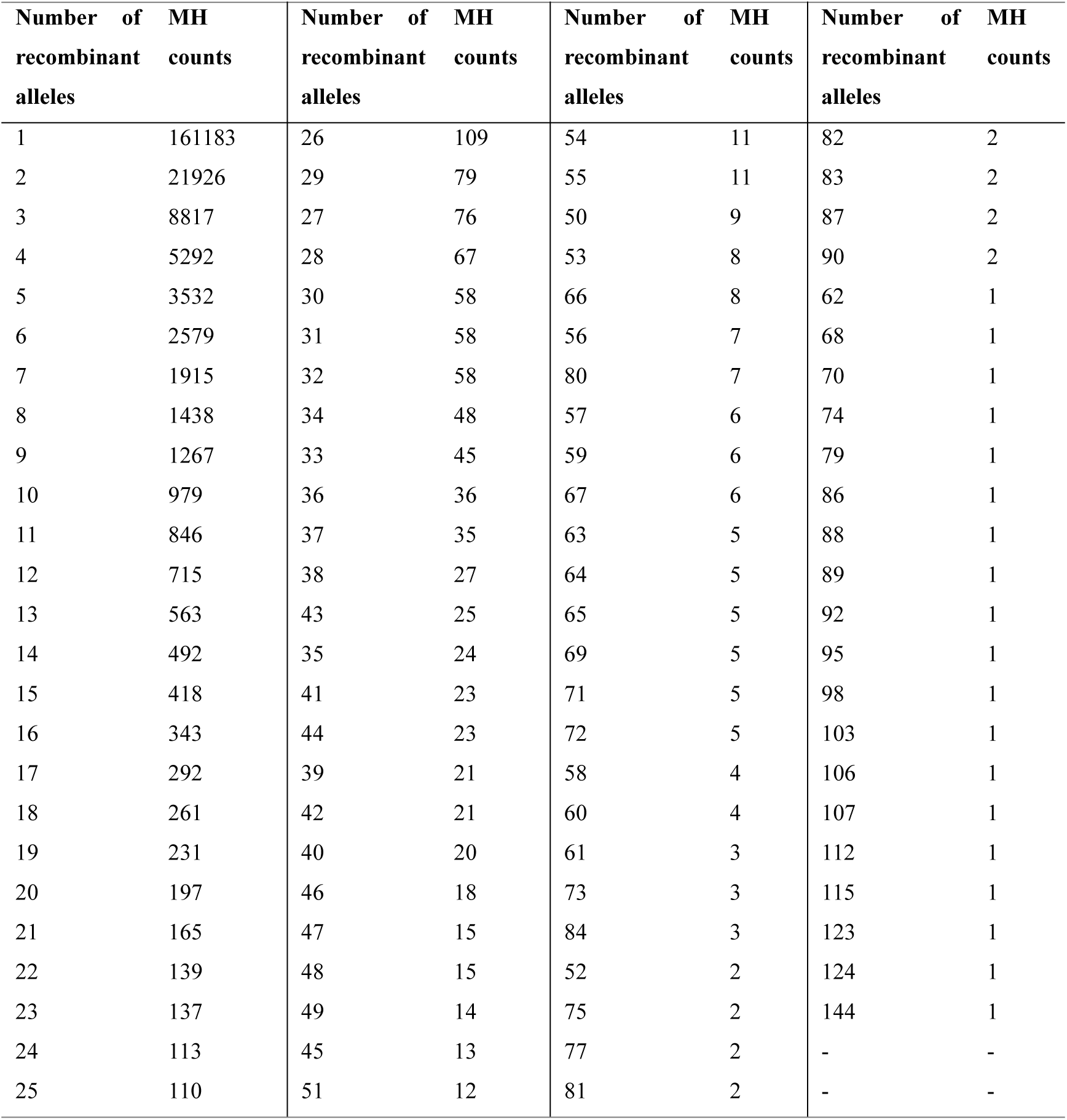
Distribution of the number of recombinant alleles in the 18-sum population.

### 3.4. The proportion of recombinant MH loci

The proportion of recombinant MH loci (Pro_r) of each population ranged from 0.02% (ITU) to 0.69% (GWD) (Fig. 7A) and increased with the increase of Ae value (Fig. 8A). Five 1000 Genomes Project super populations (AFR: African, AMR: Admixed American, EAS: East Asian, EUR: European, SAS: South Asian) possessed different proportions of recombinant MHs. There was a significant difference in *Pro*_r between the super populations AFR and SAS (p-value < 0.05), but not between AFR and the other three super populations. When Ae≥4 (including 4–5 and ≥5), CHS had a higher proportion of recombinant MHs than others relatively, while ITU had the least. With Ae≥5, CHS had 55.17% recombinant loci and ITU had 6.25% (Fig. 8A). Moreover, the recombination occurred in all MHs with Ae≥8 (Table 3). The Pearson correlation coefficient between Ae intervals and the proportion of recombinant MH loci was 0.72 (Fig. 8D).

**Figure 7.**
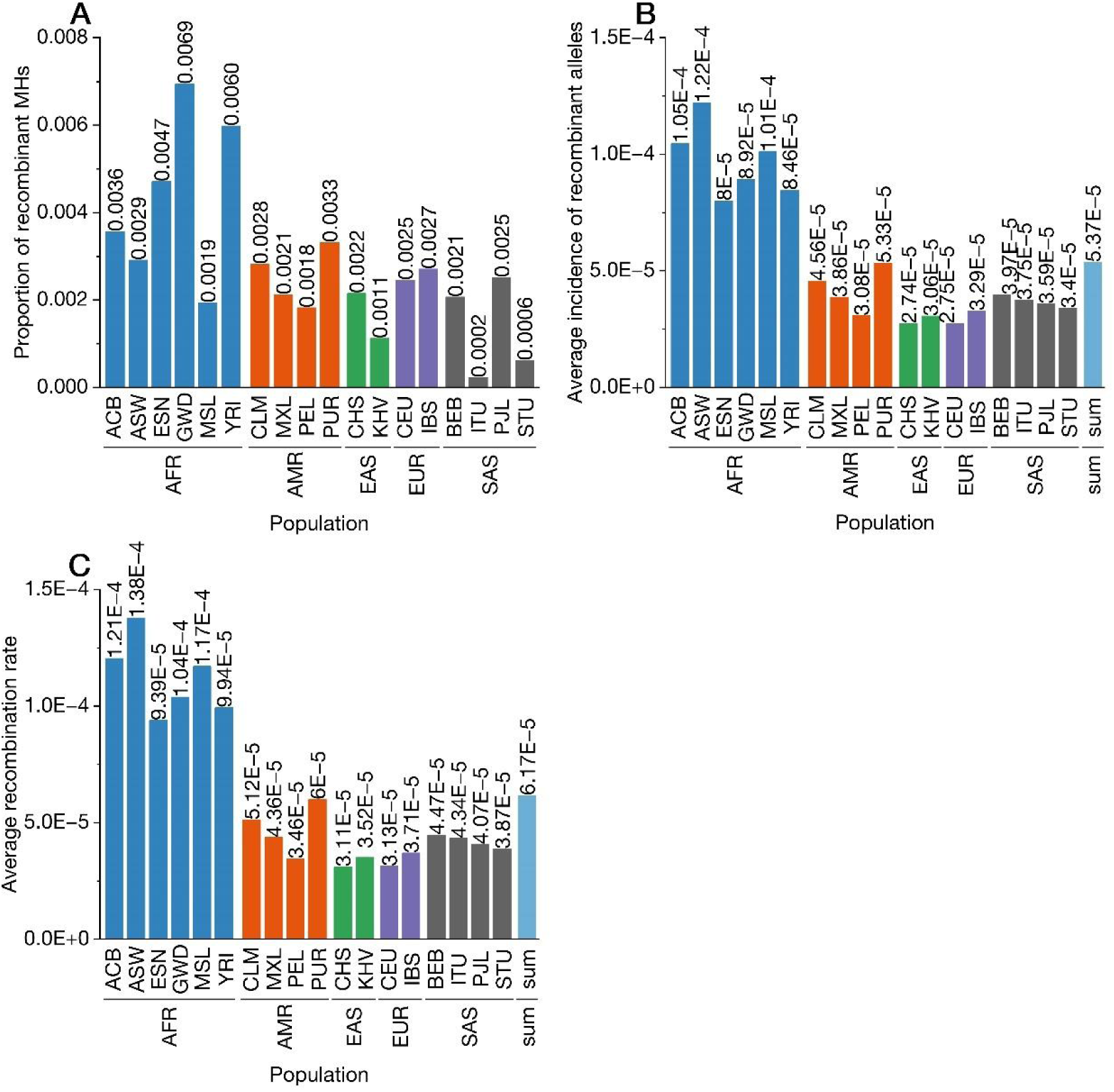
Pro_r, AvgP, and AvgRR for both the sub and super populations. A: The proportion of recombinant MH loci (Pro_r) of each population. B: The average incidence of recombinant alleles (AvgP) of each population and the 18-sum population. C: The average recombination rate (AvgRR) of each population and the 18-sum population. Each color represents a super population except for the 18-sum.

**Figure 8.**
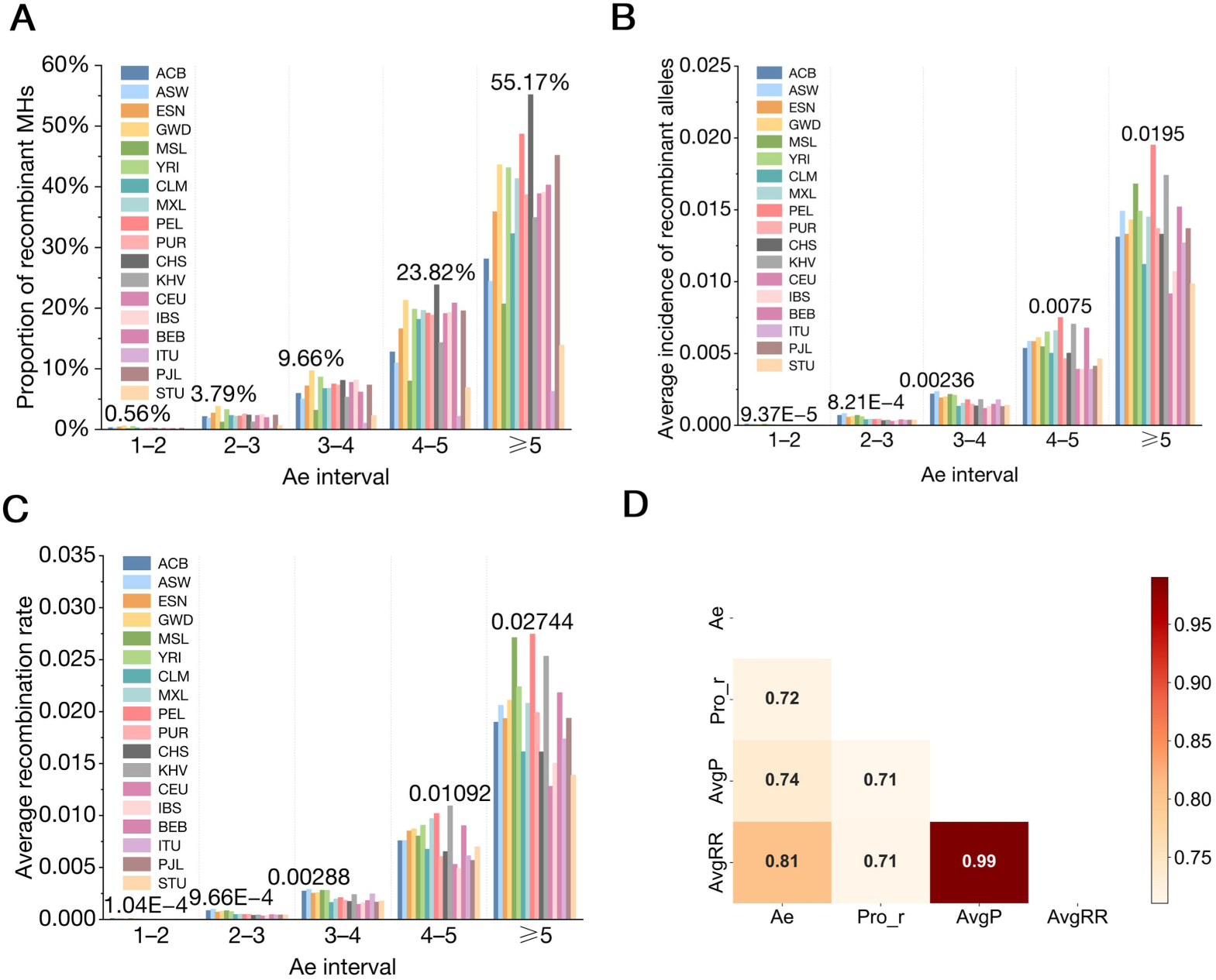
Relationship between three different measures and Ae intervals (1–2, 2–3, 3–4, 4–5, ≥5) for each population. A: Relationship between the proportion of recombinant MHs (Pro_r) and Ae value for each population. B: Relationship between the average incidence of recombinant alleles (AvgP) and Ae value for each population. C: Relationship between the average recombination rate (AvgRR) and Ae value for each population. In the Ae interval of 1–2, 2–3, 3–4, 4–5 and ≥5, the populations with the highest recombination rate are ASW, ASW, ASW, KHV, and PEL, respectively. The values are marked on the specific bars. D: Pearson correlation coefficients between three different measures and Ae interval.

**Table 3.** Information on the proportion of recombinant loci, the average incidence of recombinant alleles, and the average recombination rate of 18 populations for microhaplotypes in different Ae value intervals.

| Ae | Pro_r | AvgP | AvgRR | Number of MHs |
| --- | --- | --- | --- | --- |
| 1–2 | 2.72% | 4.38E-05 | 4.95E-05 | 6,840,658 |
| 2–3 | 19.28% | 4.42E-04 | 5.33E-04 | 136,031 |
| 3–4 | 40.51% | 1.65E-03 | 2.11E-03 | 6,226 |
| 4–5 | 67.67% | 6.17E-03 | 8.87E-03 | 464 |
| 5–6 | 86.02% | 1.12E-02 | 1.62E-02 | 93 |
| 6–7 | 90.48% | 1.72E-02 | 2.37E-02 | 21 |
| 7–8 | 92.86% | 1.87E-02 | 2.51E-02 | 14 |
| 8–9 | 100.00% | 3.03E-02 | 3.41E-02 | 2 |
| 9–10 | - | - | - | 0 |
| 10–11 | - | - | - | 0 |
| 11–12 | 100.00% | 6.73E-02 | 8.80E-02 | 1 |
| 12–13 | - | - | - | 0 |
| 13–14 | - | - | - | 0 |
| 14–15 | - | - | - | 0 |
| 15–16 | - | - | - | 0 |
| 16–17 | - | - | - | 0 |
| 17–18 | - | - | - | 0 |
| 18–19 | - | - | - | 0 |
| 19–20 | 100.00% | 3.57E-02 | 5.73E-02 | 1 |
| ≥20 | - | - | - | 0 |
| Total | 3.08% | 5.37E-05 | 6.17E-05 | 6,983,511 |

### 3.5. The average incidence of recombinant alleles

In the CHS population, 82.69% of recombinant MHs had only one recombinant allele, and 96.29% of recombinant MHs possessed 1–3 recombinant alleles as the Ae value fell in 1– 4 (Fig. 9), as the similar trend with the other 17 populations.

**Figure 9.**
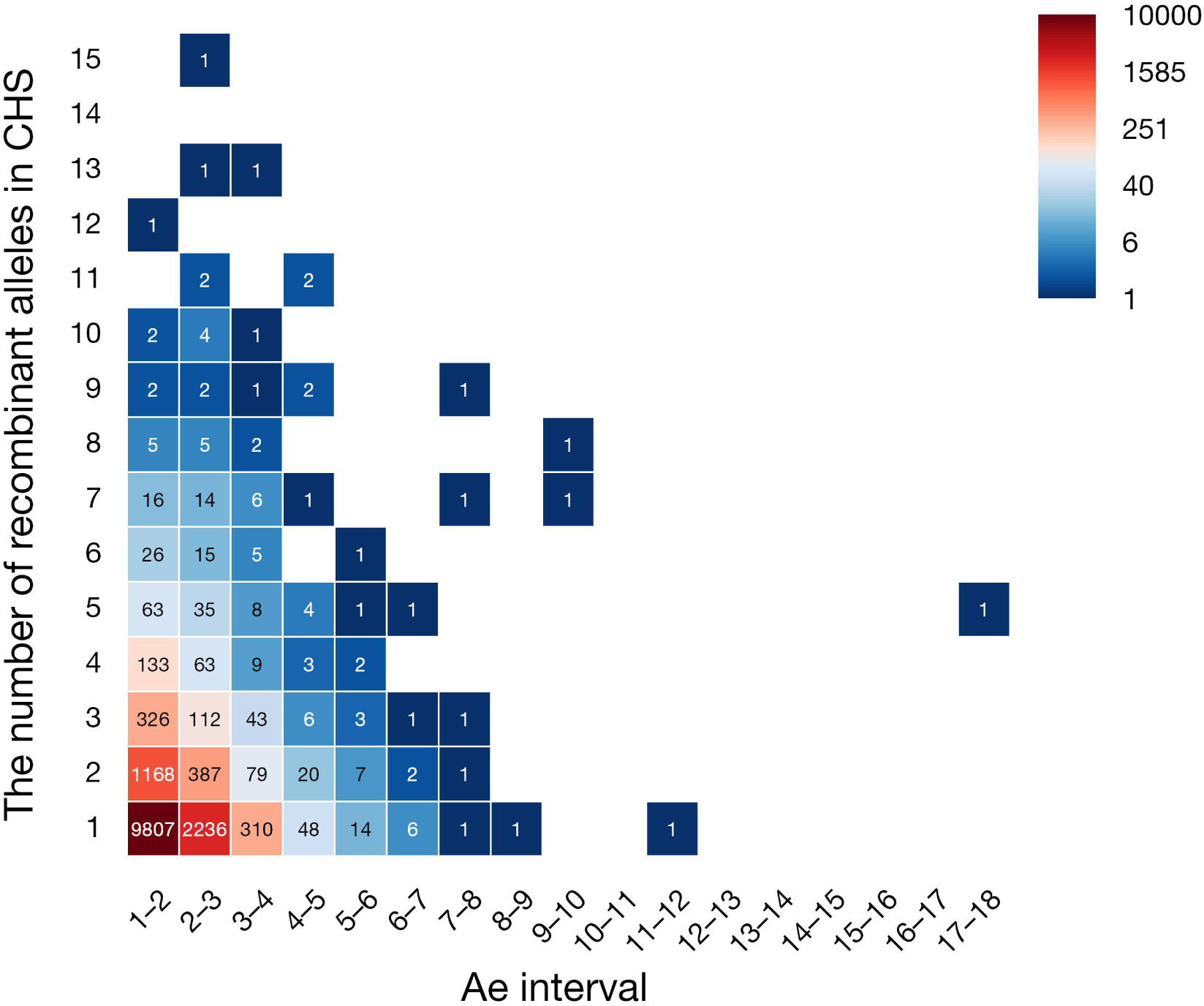
The number of MHs and the number of recombinant alleles in the CHS population based on different Ae intervals. Data value annotation in each cell represents the number of MHs.

Of the 18 populations, the ASW population obtained the highest average incidence of recombinant alleles at 1.22 ⊆ 10^-4^ meaning that the probability of producing recombinant alleles in 6,983,511 MHs for 13 trios of ASW was 0.0122% while it was 5.37⊆10^-5^ for the 18-sum population (Fig. 7B). AvgP of the super population AFR was higher than that of other four super populations (p-value < 0.05). There is a clear trend of increasing for the average incidence of recombinant alleles rising with the growth of Ae for each population (Fig. 8B) and 18-sum population (Table 3). The probabilities of producing recombinant alleles are 0.0044% with Ae between 1–2, 0.044% with Ae between 2–3, 0.17% with Ae between 3–4, 1%–2% with Ae between 5–8, 3%–4% with Ae 8–9 and 19–20. Especially, AvgP was 6.73% between Ae 11–12. The Pearson correlation coefficient between the Ae interval and the average incidence of recombinant alleles was 0.74 (Fig. 8D), indicating a strong relationship between the two variables.

### 3.6. The average recombination rate of MHs

We counted the number of recombinations within MHs of 1204 meioses and calculated the average recombination rate (AvgRR) for each population ranging from 10^-5^ to 10^-4^ (Fig. 8C, Table 4) with an average of 6.17 ⊆ 10^-5^ as a whole (Table 3). In other words, the probability of recombination occurring within MHs was approximately ⊆ 10^-5^ among 18 populations. With the Ae value increasing, the average recombination rate increased rapidly. While the average recombination rate of MHs with Ae 1–2 was 4.95⊆10^-5^ less than that of all MHs, the rate increased to ⊆10^-3^ for MHs with Ae 3–4 and ⊆10^-2^ for MHs with Ae≥5 and peaked at Ae 11–12: 8.80 ⊆ 10^-2^ (Table 3). Of the 18 populations, ASW had the highest average recombination rate at 1.38×10^-4^ with CHS having the lowest average recombination rate at 3.11×10^-5^ (Fig. 7C). Besides, the AvgRR of the super population AFR was higher than that of other four super populations (p-value < 0.05). Accordingly, the Pearson correlation coefficient between Ae intervals and the average recombination rate for the 18-sum population was 0.81 (Fig. 8D).

**Table 4.**
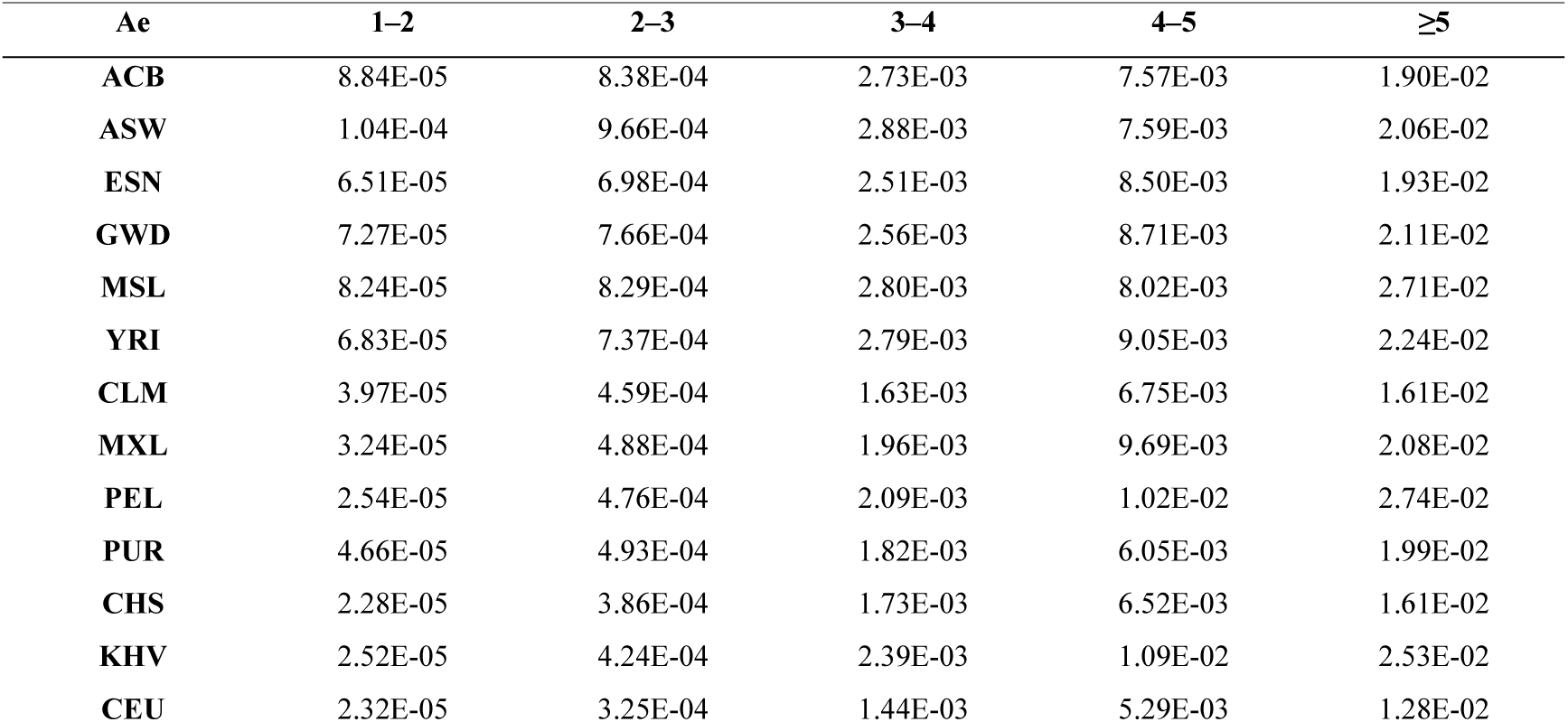
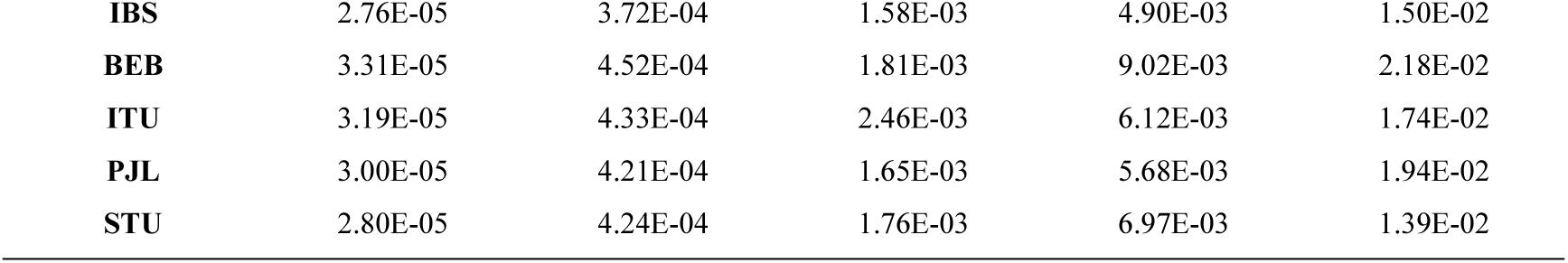
The average recombination rate of 18 populations with different Ae intervals.

| Ae | 1–2 | 2–3 | 3–4 | 4–5 | $\geq 5$ |
| --- | --- | --- | --- | --- | --- |
| ACB | 8.84E-05 | 8.38E-04 | 2.73E-03 | 7.57E-03 | 1.90E-02 |
| ASW | 1.04E-04 | 9.66E-04 | 2.88E-03 | 7.59E-03 | 2.06E-02 |
| ESN | 6.51E-05 | 6.98E-04 | 2.51E-03 | 8.50E-03 | 1.93E-02 |
| GWD | 7.27E-05 | 7.66E-04 | 2.56E-03 | 8.71E-03 | 2.11E-02 |
| MSL | 8.24E-05 | 8.29E-04 | 2.80E-03 | 8.02E-03 | 2.71E-02 |
| YRI | 6.83E-05 | 7.37E-04 | 2.79E-03 | 9.05E-03 | 2.24E-02 |
| CLM | 3.97E-05 | 4.59E-04 | 1.63E-03 | 6.75E-03 | 1.61E-02 |
| MXL | 3.24E-05 | 4.88E-04 | 1.96E-03 | 9.69E-03 | 2.08E-02 |
| PEL | 2.54E-05 | 4.76E-04 | 2.09E-03 | 1.02E-02 | 2.74E-02 |
| PUR | 4.66E-05 | 4.93E-04 | 1.82E-03 | 6.05E-03 | 1.99E-02 |
| CHS | 2.28E-05 | 3.86E-04 | 1.73E-03 | 6.52E-03 | 1.61E-02 |
| KHV | 2.52E-05 | 4.24E-04 | 2.39E-03 | 1.09E-02 | 2.53E-02 |
| CEU | 2.32E-05 | 3.25E-04 | 1.44E-03 | 5.29E-03 | 1.28E-02 |
| **IBS** | 2.76E-05 | 3.72E-04 | 1.58E-03 | 4.90E-03 | 1.50E-02 |
| **BEB** | 3.31E-05 | 4.52E-04 | 1.81E-03 | 9.02E-03 | 2.18E-02 |
| **ITU** | 3.19E-05 | 4.33E-04 | 2.46E-03 | 6.12E-03 | 1.74E-02 |
| **PJL** | 3.00E-05 | 4.21E-04 | 1.65E-03 | 5.68E-03 | 1.94E-02 |
| **STU** | 2.80E-05 | 4.24E-04 | 1.76E-03 | 6.97E-03 | 1.39E-02 |

### 3.7. Referenced recombination rate based on HapMap

We used referenced data of HapMap to obtain the referenced AvgRR of MHs to compare with AvgRR produced by our trios-based observation. After the physical coordinates of the HapMap genetic map had been converted to GRCh38, 93.34% of MHs acquired the referenced rate of recombination (ld_based_rate), the rest encompassing loci lack of the information of the genetic map and those of the genetic map being zero. The recombinant MHs with valid ld_based_rate values queried were divided into 5 groups by ld_based_rate. We counted these MH loci in each ld_based_rate interval and calculated the average recombination rate of MH loci in the corresponding interval (Fig. 10A). Under the unit of cM/Mb based on HapMap, if the ld_based_rate of the locus is 1 cM/Mb, the average referenced rate of recombination within the locus is about 10^-6^ according to the estimated value that 1 cM equals to 1% recombination rate. In essence, both average recombination rates acquired from HapMap and our study represented the average level of recombination rate of DNA sequence covered by MHs. When the ld_based_rate of MHs was higher than 10 cM/Mb, the AvgRR of our study, 10^-6^ – 10^-5^, was essentially in line with it, while AvgRR was higher than ld_based_rate when it was lower than 10 cM/Mb. The average recombination rate of all MHs with ld_based_rate≥1 cM/Mb was about 1.99 ⊆ 10^-5^ while that of all MHs with ld_based_rate<1 cM/Mb was about 1.35⊆10^-5^ (Fig. 10D).

**Figure 10.**
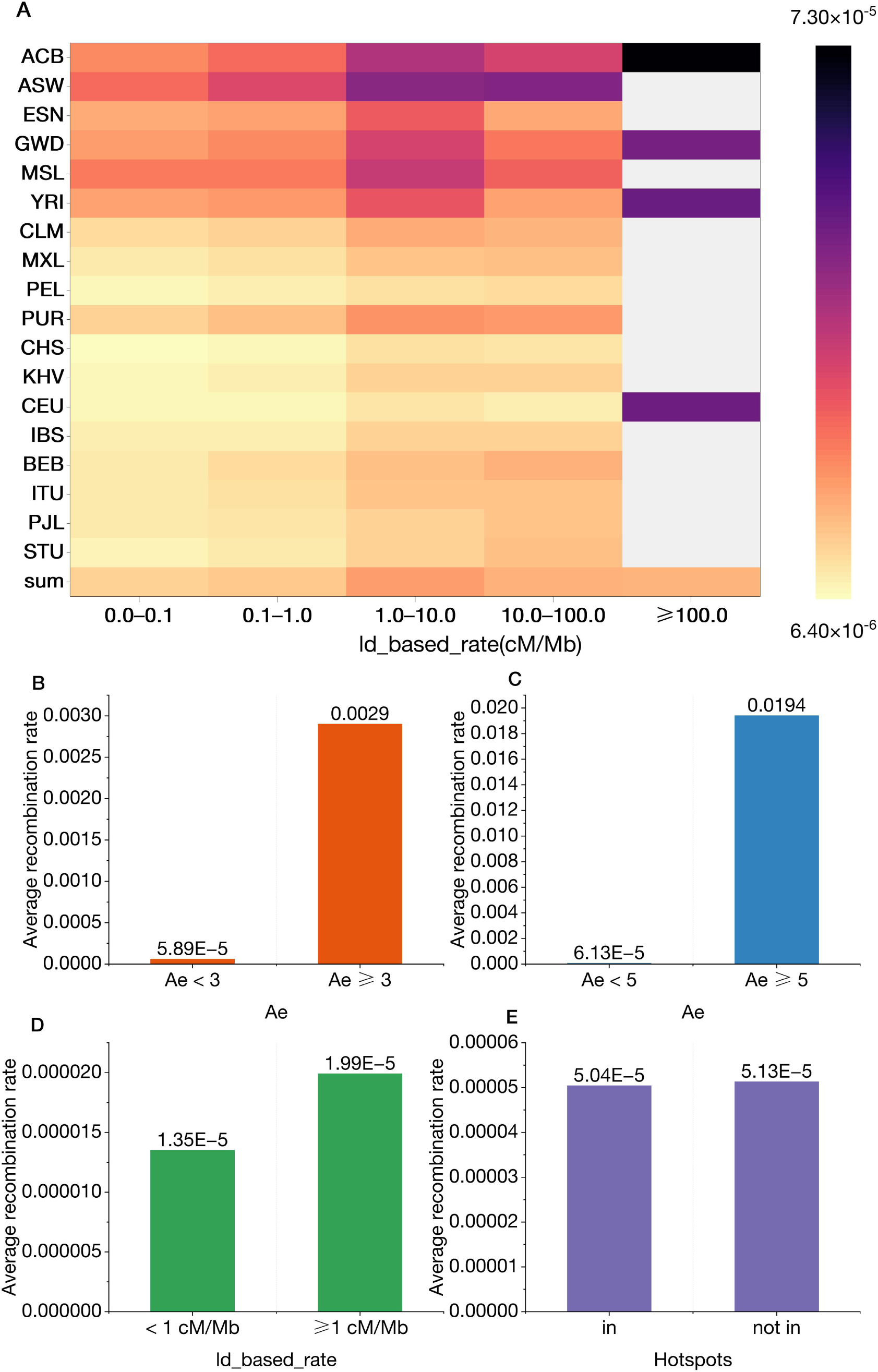
Influences of different factors on AvgRR. A: The average recombination rate of MHs in the corresponding ld_based_rate interval in the 18 and 18-sum populations. B, C, D, E: Effect of Ae value, ld_based_rate, and recombination hotspots on the average recombination rate of MHs, respectively.

Referring to 32,996 recombination hotspots where 18 hotspots had been deleted during conversion of coordinates published by HapMap (build 35), 14,473 recombinant MHs accounting for 7.21% of all recombinant MHs distributed in recombination hotspots, as 186,264 recombinant MHs accounting for 92.79% located the outside of recombination hotspots. The rest 14,314 recombinant MHs failed to get ld_based_rate, or it was 0. The average recombination rates were 5.04⊆10^-5^ and 5.13⊆10^-5^ for all MHs distributing inside and outside the recombination hotspots, respectively (Fig. 10E).

### 3.8. Removing partial high-density distribution regions

We observed some high-density distribution regions of recombinant MHs in the genome, which were mostly located near the gaps of GRCh38 (https://genome.ucsc.edu/cgi-bin/hgTables?db=hg38&hgta_group=map&hgta_track=gap&hgta_table=gap&hgta_doSchema=describe+table+schema) and other regions including HLA-DRB1, IL1B, 2q14.1, 2q21.2 and 3p12.3. After removing these regions and their vicinities, we retained 5,068,022 MHs, of which there were 118,936 recombinant MHs (see Fig. 11A for their distribution on the genome). The results showed that the proportion of recombinant MHs (Pro_r) with Ae 1–5 had decreased (p-value<0.0001) while the difference was not significant when Ae ≥5 (p-value>0.05) (Fig. 11B). In addition, the average incidence of recombinant alleles (AvgP) and the average of recombination rate (AvgRR) both had significantly decreased in the same Ae interval (p-value<0.0001) among the 18 populations (Fig. 11C, D). Taking the 18 populations as a whole, we observed a reduction in AvgP and AvgRR between Ae 1–8 and a reduction in Pro_r between Ae 1–5. Pro_r, AvgP, and AvgRR were overall lower (Table 5).

**Figure 11.**
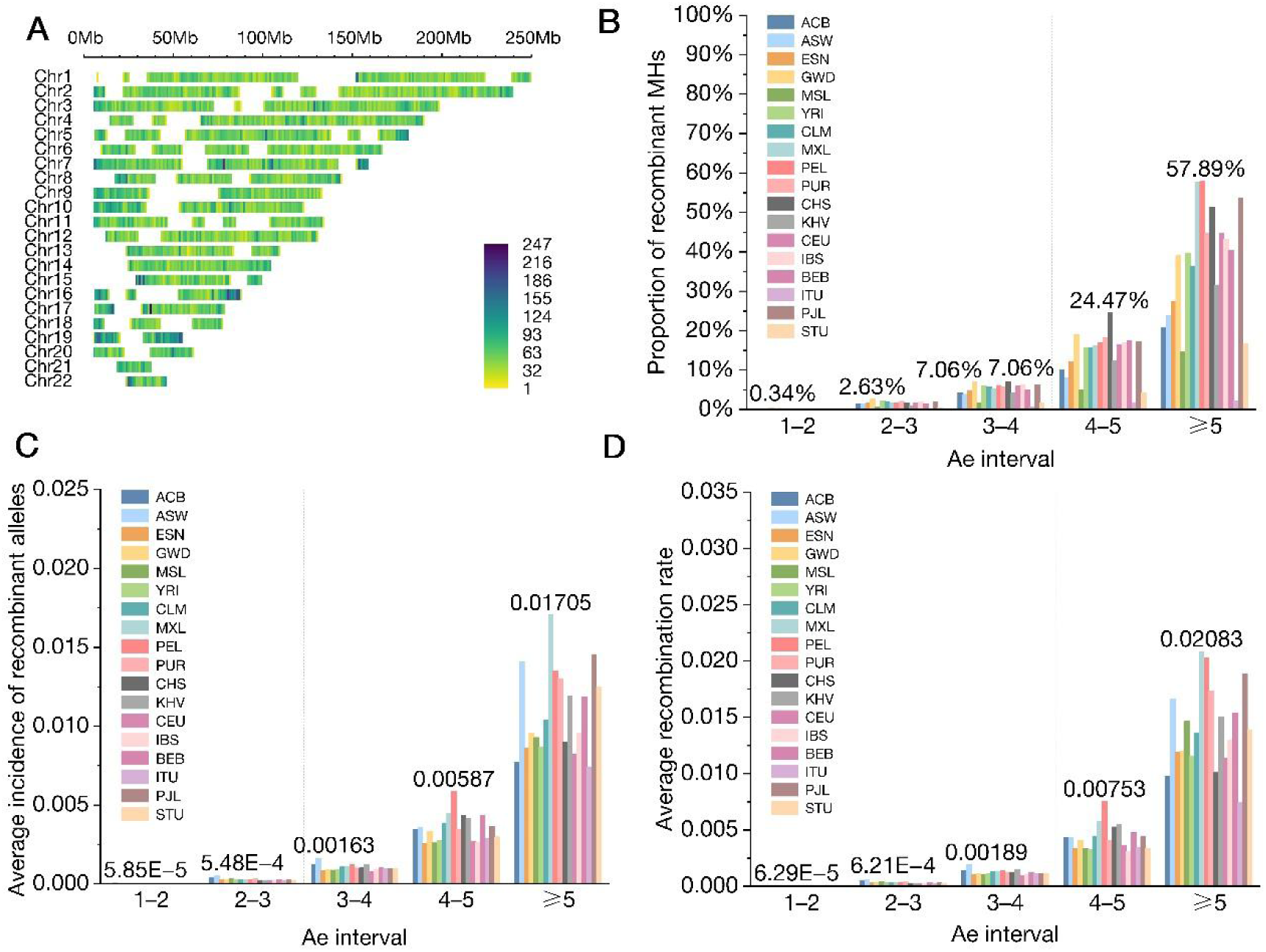
Density plot and three measures after removing partial high-density distribution regions. A: Distribution of recombinant MHs in the human genome after the deletion. B, C, D: Pro_r, AvgP, AvgRR with different Ae values, respectively.

**Table 5.** Pro_r, AvgP, and AvgRR of the 18-sum population after removing partial high-density distribution regions.

| Ae | Pro_r | AvgP | AvgRR | Number of MHs |
| --- | --- | --- | --- | --- |
| 1-2 | 2.04% | 2.31E-05 | 2.51E-05 | 4,970,646 |
| 2-3 | 16.99% | 2.87E-04 | 3.29E-04 | 93,348 |
| 3-4 | 36.56% | 1.17E-03 | 1.41E-03 | 3,777 |
| 4-5 | 66.67% | 4.50E-03 | 5.57E-03 | 213 |
| 5-6 | 93.10% | 1.13E-02 | 1.56E-02 | 29 |
| 6-7 | 100.00% | 1.18E-02 | 1.60E-02 | 8 |
| 7-8 | - | - | - | 0 |
| 8-9 | 100.00% | 5.90E-02 | 6.64E-02 | 1 |
| ≥9 | - | - | - | 0 |
| Total | 2.35% | 2.91E-05 | 3.21E-05 | 5,068,022 |

## 4. Discussion

We studied the recombination within MHs by directly observing the inheritance between parental and offspring’s MH alleles on the basis of trios. This intuitive analysis method to the resolution rate of recombination events is mainly ascribed to two factors: the density of genetic markers detected and the number of meioses observed. Unlike most research based on the family method, our work on the resolution rate of recombination events is hardly limited by the density of genetic markers. The precision of the early family method capturing recombination is 10 kb approximately [14] while it is 350 bp in our study. However, about 23% of a genome sequence uncovered in the MH library is not involved in the work, and recombination between (not within) MHs remained to be observed. Our results substantiated that MH alleles of offspring were not always identical to one of the paternal or maternal alleles during inheritance, within which recombination events occurred more widely and frequently than expected. The average recombination rate of MHs was approximately 6.17 ⊆ 10^-5^, which was between the mutation rate of SNP (10^-8^) and STR (10^-3^). After eliminating the influences of sequencing error and chromosomal structure, recombination within MHs was still a force to be reckoned with. We removed partial sequencing regions with high-density recombinant MHs, which might be caused by unknown sequences [20] such as pericentromeric, subtelomeric regions, and “falsely duplicated and poorly assembled sequences” [21], et al. The sequence gaps were flanked by duplications, regions of which were sites of genetic instability and large-scale structural polymorphism, complicating their sequence and assembly [22]. HLA-DRB1 has abundant polymorphism to affect the recombination observed [23, 24]. Pro_r, AvgP, and AvgRR were overall lower than before but remained in the same order of magnitude, evincing that the influences were limited and the results were reliable and universal.

The 18 populations from 5 continents accentuated the discrepancy of recombination within MHs between continents. Particularly, the number of recombinant alleles and the number of recombinant MHs in the AFR were much higher than that in the other four continents, illustrating the more active recombination events in Africa. This implied that the faster decay of linkage disequilibrium length with physical distance in Africa at the population level was not only due to differences in population history [25], but might also be related to the more even distribution of locations of DNA recombination in African than in other continents. The analogous study [26] found that on a finer scale of less than 3 Mb, the genetic map of the European population differed significantly from the YRI population and African Americans with more recombination hotspots detectable in West Africa. Of 215,051 recombinant MHs, only 4 loci were shared by the 18 populations while the vast majority of them were “non-shared” and occurred specifically across populations. The paucity of trios in some populations could be one of contributing factors to the small number of shared loci. In addition to this, the unshared African-enriched hotspots in Africans, which has been reported before [26], suggested that different populations possessed a similar recombination rate at the genome level as the physical positions of recombination were various across populations, meaning that there were differences in recombination rates across populations at a finer scale. Meanwhile, the discrepancy might be amplified by the limited meiosis.

We further explored the correlation of Ae intervals and the proportion of recombinant MH loci (Pro_r), the average incidence of recombinant alleles (AvgP), and the average recombination rate (AvgRR) separately. It was overt that the probability of recombination within MHs increased significantly with increasing Ae value, regardless of the measure. From Ae 1–2 to Ae ≥5, the order of magnitude of AvgP or AvgRR increased from -5 to -2. Moreover, the Pearson correlation coefficients of Ae intervals and each of them were 0.72, 0.74, and 0.81, respectively, thus sustaining a strong correlation between them as well. It hinted that the mechanism of MH polymorphism might be closely related to the recombination between SNPs constituting MHs. While the average recombination rate of MHs with low Ae value was low, we still strongly recommended scrupulous selection when applying them to kinship analysis. The observation of family data before using specific MHs was suggested, which made for the filtering of MHs.

A comparison of the results between our study and HapMap recombination hotspots proved that recombinant MHs were not always concentrated in the region of hotspots. The recombination hotspot from HapMap was not a crucial factor for the average rate of MH recombination. The average recombination rates of MHs distributed outside and inside the recombination hotspots were both around 10^-5^, with a petty difference. When ld_based_rates of MHs were between 0–10 cM/Mb, the average recombination rate (AvgRR) of the 18 populations was 1 to 2 orders of magnitude higher than theirs. There were two explanations for this, including the resolution of recombination, which reached 350 bp and thereby encompassing noncrossover gene conversion [27–29], and the direct observation method, which captured recombination that occurred during gamete production, rather than inferring population recombination rates in terms of historical recombination events from population genetics methods. The population genetics methods were so conservative that only a few recombination events could be detected [30, 31]. We did not discriminate crossovers and gene conversion in this study. Noncrossover gene conversion usually occurred within only tens to thousands of base pairs [29] and did not interfere with one another as much as crossovers did [32, 33], explaining the phenomenon of multi-recombination events in a short segment. Meiotic recombination was initiated from double-strand breaks (DSBs), the majority of which did not lead to crossovers but ended as gene conversions [27], implying that a significant number of recombination events in our study most likely originated from gene conversions.

Considering the recombination events within MHs and the definition of the haplotype, it sounded more legitimate to call microhaplotypes as multi-SNPs in that these SNPs were not always inherited together. In addition to the results from 1KG, more in-house sequencing data of trios were expected to validate the reliability of the phenomenon observed. Generally, we observed the recombination rate in the genome at a small scale and suggested the appraisal of the recombination rate of candidate MH loci before choosing them to perform kinship analysis, and the calculation of the likelihood ratio involving recombination rate within MHs.

## 5. Conclusion

This study has filtered and constituted the MH set covering 77.21% of the human genome sequence on the basis of e1kGP. The results of this study indicate that MHs within which recombination has occurred have a wide distribution in the genome. In the meantime, their positions are not limited to recombination hotspots, and the population differences in quantity and distribution are observed for these loci. Also, recombination plays an important role in the mechanism of MH polymorphism, and with high Ae values, it is more likely for MH loci to generate variant haplotypes or alleles due to recombination during inheritance. It is, therefore, necessary for investigation of the recombination rate for each MH locus, or correction to probabilistic models using the average recombination rate of MHs.

## Acknowledgments

We thank Kenneth K. Kidd for providing useful recommendations for the manuscript. This work was supported by grants from the National Natural Science Foundation of China (no. 82230064, 82072120, and 81630054).

