## Supplemental Table 3 for "Evaluation of recombination rate of microhaplotypes based on the 1000 Genomes Project family data"

| Ae Interval | YRI | CLM | CHS | CEU | PJL | ACB | ASW | ESN | GWD | MSL | MXL | PEL | PUR | KHV | IBS | BEB | ITU | STU |
| --- | --- | --- | --- | --- | --- | --- | --- | --- | --- | --- | --- | --- | --- | --- | --- | --- | --- | --- |
| 1–2 | 6,745,266 | 6,828,679 | 6,855,153 | 6,837,691 | 6,831,651 | 6,748,808 | 6,760,486 | 6,746,686 | 6,745,336 | 6,739,484 | 6,846,793 | 6,867,930 | 6,815,940 | 6,856,064 | 6,832,914 | 6,833,725 | 6,836,606 | 6,834,765 |
| 2–3 | 219,972 | 146,497 | 122,158 | 137,915 | 143,483 | 217,219 | 206,952 | 218,599 | 220,000 | 224,530 | 129,813 | 110,760 | 158,176 | 121,042 | 142,333 | 141,562 | 138,566 | 140,467 |
| 3–4 | 16,010 | 7,568 | 5,752 | 7,242 | 7,618 | 15,408 | 14,191 | 16,039 | 15,951 | 17,050 | 6,298 | 4,484 | 8,524 | 5,880 | 7,511 | 7,504 | 7,596 | 7,547 |
| 4–5 | 1,776 | 612 | 361 | 524 | 604 | 1,599 | 1,460 | 1,713 | 1,715 | 1,944 | 474 | 261 | 690 | 399 | 602 | 571 | 599 | 595 |
| 5–6 | 335 | 103 | 60 | 101 | 98 | 341 | 293 | 322 | 335 | 323 | 93 | 54 | 122 | 81 | 95 | 104 | 98 | 93 |
| 6–7 | 92 | 32 | 16 | 23 | 36 | 77 | 78 | 97 | 106 | 108 | 24 | 12 | 34 | 32 | 32 | 29 | 24 | 24 |
| 7–8 | 31 | 10 | 6 | 9 | 15 | 29 | 26 | 32 | 34 | 35 | 10 | 7 | 14 | 7 | 15 | 8 | 16 | 11 |
| 8–9 | 10 | 6 | 1 | 1 | 2 | 13 | 13 | 9 | 20 | 21 | 2 | 2 | 5 | 3 | 4 | 6 | 3 | 5 |
| 9–10 | 11 | 1 | 2 | 2 | 2 | 9 | 7 | 8 | 5 | 5 | 2 | 1 | 2 | 2 | 3 | 0 | 1 | 2 |
| 10–11 | 5 | 1 | 0 | 1 | 0 | 5 | 3 | 2 | 5 | 4 | 1 | 0 | 2 | 0 | 1 | 1 | 1 | 0 |
| 11–12 | 0 | 1 | 1 | 0 | 0 | 1 | 1 | 1 | 2 | 5 | 0 | 0 | 0 | 0 | 0 | 0 | 0 | 0 |
| 12–13 | 1 | 0 | 0 | 0 | 0 | 0 | 0 | 0 | 0 | 0 | 0 | 0 | 0 | 0 | 0 | 0 | 0 | 0 |
| 13–14 | 1 | 1 | 0 | 0 | 0 | 0 | 0 | 2 | 1 | 1 | 1 | 0 | 0 | 0 | 0 | 0 | 0 | 1 |
| 14–15 | 0 | 0 | 0 | 0 | 1 | 1 | 0 | 0 | 0 | 0 | 0 | 0 | 0 | 0 | 0 | 0 | 0 | 0 |
| 15–16 | 1 | 0 | 0 | 0 | 0 | 0 | 0 | 0 | 1 | 0 | 0 | 0 | 0 | 0 | 0 | 0 | 0 | 0 |
| 16–17 | 0 | 0 | 0 | 0 | 0 | 0 | 0 | 0 | 0 | 0 | 0 | 0 | 1 | 0 | 0 | 0 | 0 | 0 |
| 17–18 | 0 | 0 | 1 | 1 | 0 | 0 | 0 | 1 | 0 | 0 | 0 | 0 | 0 | 0 | 0 | 1 | 0 | 0 |
| 18–19 | 0 | 0 | 0 | 0 | 0 | 0 | 0 | 0 | 0 | 0 | 0 | 0 | 0 | 0 | 0 | 0 | 0 | 0 |
| 19–20 | 0 | 0 | 0 | 0 | 0 | 0 | 0 | 0 | 0 | 0 | 0 | 0 | 1 | 0 | 0 | 0 | 0 | 0 |
| 20–21 | 0 | 0 | 0 | 0 | 0 | 0 | 0 | 0 | 0 | 0 | 0 | 0 | 0 | 1 | 0 | 0 | 1 | 0 |
| 21–22 | 0 | 0 | 0 | 0 | 0 | 0 | 0 | 0 | 0 | 0 | 0 | 0 | 0 | 0 | 0 | 0 | 0 | 0 |
| 22–23 | 0 | 0 | 0 | 0 | 1 | 0 | 0 | 0 | 0 | 1 | 0 | 0 | 0 | 0 | 0 | 0 | 0 | 0 |
| 23–25 | 0 | 0 | 0 | 1 | 0 | 0 | 1 | 0 | 0 | 0 | 0 | 0 | 0 | 0 | 0 | 0 | 0 | 0 |
| ≥25 | 0 | 0 | 0 | 0 | 0 | 1 | 0 | 0 | 0 | 0 | 0 | 0 | 0 | 0 | 1 | 0 | 0 | 1 |

**Supplemental Table S3**. Distribution of Ae of MH loci in different populations.
